# The Impact of Clinical and Molecular Variant Properties on Calibration and Performance of Variant Effect Prediction Tools

**DOI:** 10.1101/2024.10.02.614907

**Authors:** Ofer Isakov, Reut Fluss, Dina Marek-Yagel, Shay Ben Shachar

## Abstract

**Background:** Variant Effect Prediction (VEP) tools are essential for determining the potential pathogenicity of genetic variants, aiding clinical diagnostics and genetic counseling. However, their performance can vary depending on molecular and clinical contexts, complicating variant classification.

**Aim:** This study aims to assess the performance variability of commonly used VEP tools under different conditions. Additionally, the study aims to recalibrate score thresholds to better reflect evidence of pathogenicity.

**Methods:** ClinVar variants classified as pathogenic (P), likely pathogenic (LP), benign (B), or likely benign (LB) were analyzed using 25 VEP tools. Tools were evaluated based on discriminatory performance. Data were stratified by variant creation date, allele frequency, conservation level, mode of inheritance (MOI), and disease category. For each subset, Bayesian methods were employed to recalibrate score thresholds corresponding to the levels of evidence defined by the American College of Medical Genetics (ACMG).

**Results:** The performance of VEP tools varied significantly across different subsets. Variants created after 2020 showed a mild yet significant decrease in the performance of certain VEP tools, particularly those trained on earlier datasets. VEP tools exhibited reduced accuracy for variants with higher allele frequencies, particularly those exceeding a frequency of 10^−4^, suggesting a need for recalibration when assessing more common variants. Tools demonstrated lower discriminatory performance for variants located in regions with high conservation, mostly due to a decrease in specificity. Variants affecting autosomal recessive (AR) and X-linked (XL) genes were more accurately classified compared to those affecting autosomal dominant (AD) genes by most tools. Differences in MOI and conservation levels within certain disease categories were shown to correlate with overall performance. Recalibration of prediction scores resulted in lower score thresholds in low conservation regions compared to high and uncovered subsets in which higher levels of evidence may be achieved.

**Conclusion:** VEP tools exhibit context-dependent performance variability, necessitating score recalibration for accurate classification.

## BACKGROUND

Variant Effect Prediction (VEP) tools are computational methods designed to assess the potential impact of genetic variants on gene function. These tools are mainly utilized to distinguish between benign variants, which do not cause disease, and pathogenic variants, which are associated with disease phenotypes. This distinction is pivotal for clinical diagnostics, genetic counseling, and in genetic disorders research. Several pathogenicity prediction tools have been developed, each utilizing different algorithms and datasets to predict the functional impact of variants. While early prediction tools primarily relied on inter-species conservation and limited functional data, newer tools now incorporate a plethora of characteristics, including protein structure, variant frequency in primates and the scores of previously developed models, as additional features to improve performance. The underlying algorithms also vary significantly across prediction tools, with applications of regression models, machine learning methods, model ensembles and deep neural networks. Even with the advent of these tools, variant classification remains challenging and the analytic burden of variants of unknown significance (VUS) hinders the task of automating this task in clinical settings.

Variant classification guidelines published in 2015 by the American College of Medical Genetics and Association for Molecular Pathology (ACMG/AMP) recommend the use of in-silico prediction tools to modify the probability for pathogenicity of missense variants^1^. In these guidelines, predictor evidence was limited to supporting strength due to the relatively low sensitivity and specificity of the predictors available at that time. Since 2015, an abundance of prediction tools have been published and compared against existing ones, usually demonstrating improved performance across different curated data sets such as ClinVar^2^, gnomAD^3^ and multiplexed assays of variant effect (MAVE). In a recent publication by the ClinGen Sequence Variant Interpretation (SVI) Working Group, the performance and calibration of some of these tools has been re-assessed, and new thresholds were corresponding to stronger evidence levels compared to the original 2015 guidelines^4^.

As more tools are developed with a higher amount of features and tested using subsets of similar data sets, the possibility of inflated performance results and overfitting increases. Moreover, studies have shown that the performance of available tools can vary significantly depending on the type and context of the variant being analyzed^5–8^. This variability can lead to inconsistent predictions, complicating the clinical classification of variants. A recent study, reviewing the performance of two high performing tools (REVEL and BayesDel) per gene, has also shown that even after implementing the genome-wide-based calibration suggested by the SVI, performance remains highly discordant in many genes^7^. While providing a per-gene assessment of the performance within each threshold interval, with a conclusion regarding the usability of each interval, the aforementioned study did not provide re-calibrated score thresholds that may be used instead.

While many studies have been published focusing on overall prediction performance, a comprehensive characterization in clinically meaningful subsets has yet to be performed. In this study, we evaluate and compare the performance of common pathogenicity prediction tools under various conditions in order to uncover common biases and conditions in which these tools excel and struggle with. Furthermore, for each condition, we re-calibrate the scores provided by these tools and provide modified score thresholds corresponding to each condition.

## METHODS

### Data collection

ClinVar VCF data was retrieved from the NCBI ftp on 02/05/2024^2^. Only single nucleotide variants classified (according to the CLNSIG info value) as pathogenic (P), likely pathogenic (LP), benign (B) and likely benign (LB) were selected. Variants with an ambiguous ID or without assertion criteria were also removed from all downstream analysis. Variant annotation was done using VarSeq™ v2.6.1 (Golden Helix, Inc., Bozeman, MT, www.goldenhelix.com)^9^. Prediction scores were retrieved from dbNSFP v4.5a^10^ for 25 prediction tools: SIFT^11^, SIFT4G, PolyPhen2 (HVAR/HDIV)^12^, MutationTaster^13^, FATHMM, fathmmMKL^14^, PROVEAN^15^, VEST4^16^, MetaSVM^17^, MetaLR^17^, MetaRNN^18^, REVEL^19^, GMVP^20^, PrimateAI^21^, PrimateAI3D^22^, DEOGEN2^23^, BayesDel with/without AF^24^, ClinPred^25^, VARITY^26^, AlphaMissense^27^, DANN^28^, Genocanyon^29^ and CADD^30^. Variants without any annotation in dbNSFP were excluded from downstream analysis as well.

### Data analysis

Unless specified otherwise, inter-tools comparisons were performed on a subset of the ClinVar dataset that includes only variants that were created in the database after 01/01/2023. This was done in order to minimize the number of variants in the evaluation set that were included in the training process of the various tools. Additionally, to make sure that comparison is held on the same variant set, only variants that had information for all the compared tools were included in the comparison. Discriminatory performance was evaluated and ranked using the area under the ROC curve (AUROC), sensitivity and specificity.

### Per-dataset score threshold calculation

In each tested dataset, the posterior probability for each score was calculated. A Bayesian approach was used to calculate the posterior probability of a variant being pathogenic or likely pathogenic (P/LP) given a given model score. This approach utilizes an approximation of the prior probability of a variant being P/LP derived by Pejaver et al^4^ (0.0441) and accounts for dataset-specific class ratios.

(i) We first adjust the prior probability to account for the expected discrepancy between the variant subset group-specific ratio between P/LP and B/LB variants and the known population ratio (i.e `P(P/LP)`). We consider the entire ClinVar dataset of P/LP and B/LB variants as the initial population in which the approximated prior probability of pathogenicity (`P(P/LP)`) is 0.0441. For each variant subset group, we adjust this prior probability using the subset group’s ratio between P/LP and B/LB:

~~~
adjusted P(P/LP) = P(P/LP) * (P/LP)’/((P/LP)’+(B/LB)’) / (P/LP)°/((P/LP)°+(B/LB)°)
adjusted P(B/LB) = 1-adjusted P(P/LP)
~~~

Where `(P/LP)’ and (B/LB)’` are the number of P/LP and B/LB in the variant subset group and `(P/LP)° and (B/LB)°` are the number of P/LP and B/LB in our full ClinVar dataset.

(ii) We then apply Bayes’ theorem to calculate the posterior probability P(P/LP | score) of a variant being P/LP given its score in each variant subset group:

~~~
P(P/LP | score) = (P(score | P/LP) * adjusted P(P/LP)) / P(score)
~~~

Where `P(score)` is the marginal likelihood, calculated as:

~~~
P(score) = P(score | P/LP) * adjusted P(P/LP) + P(score | B/LB) * adjusted P(B/LB)
~~~

And `P(score | P/LP)` and `P(score|B/LB)` are the likelihood estimates of observing a given score given a variant is P/LP and B/LB, respectively. Likelihood estimates are calculated using a non-parametric kernel density estimation (R `density` function). The density function allows us to capture the distribution of scores within each class and variant subset, rather than just using a single point estimate (e.g., the mean score). This non-parametric approach allows the method to adapt to the unique characteristics of the data, without making restrictive assumptions about the underlying score distributions.

The resulting probabilities represent the likelihood of a variant being P/LP, given its score, the characteristics of the variant subset group, and the known population prevalence of P/LP variants.

#### Variant creation date analysis

Variant creation dates were collected from the latest ClinVar XML files. For each tool, a linear model was fitted to assess whether variant creation at or after 2020 affected their performance, measured by AUC50. A mixed-effects model was applied to evaluate the collective effect of creation date on the performance of the top-performing tools. Linear models were fitted for each tool in order to compare performance before and after each tool’s respective training years. A literature review was performed to elucidate the ClinVar release date used during each tool’s training. Tools that were not trained on ClinVar variants or with a creation date prior to 2013 were excluded from the analysis.

#### Conservation analysis

Conservation was defined using the phyloP score based on the multiple alignments of 100 vertebrate species (phyloP100way)^31^. All ClinVar variants phyloP scores were collected and divided into three equal bins and categorized into low (phyloP<1.646), intermediate (phyloP 1.646-6.267) and high (phyloP>6.267) conservation levels accordingly. Overall effect of conservation was evaluated by applying mixed effects models for AUROC, sensitivity and specificity. Mismatch constraint information was calculated from the mismatch observed versus expected rates retrieved from gnomAD v4.1^3^. Overall effect of mismatch constraint was evaluated by applying mixed effects models for AUROC. To identify tools that perform worse in high or low constraint regions, for each tool, the mean AUROC was calculated across constraint levels, and the difference between the minimal AUROC and the mean was collected.

#### Allele Frequency analysis

Variant allele frequency was defined according to the variant frequency across ancestries (i.e joint frequency) in the gnomAD v4.1 database^3^. Allele frequencies were categorized into bins and the AF of variants without a record in gnomAD were categorized as AF=0. Overall effect of allele frequency on performance was evaluated by applying mixed effects models with AUROC as the dependent variable. The association between AF and AUROC for each tool was calculated using a linear regression with the median AF value in each bin as the independent variable.

#### Mode of inheritance analysis

Mode of inheritance for each gene was defined using information retrieved from the Gene Curation Coalition (GenCC)^32^. Only autosomal dominant (AD), autosomal recessive (AR) and X-linked (XL) modes of inheritance were considered for downstream analysis. Variants with other modes of inheritance, or for which there is more than one possible mode of inheritance, were excluded from the analysis.

#### Disease category analysis

Gene-disease associations were derived from the panels developed by PanelApp^33^. For each panel, only genes categorized as green and amber were selected, corresponding to strong and intermediate gene-disease association. For each panel, all the variants affecting any gene in that panel were marked as associated with the panel. Only panels with at least 200 P/LP and 200 B/LB variants were analyzed (N=62). For each panel, overall performance was calculated by collecting the median AUROC for each tool in each panel, and the mean of the AUROC in all other panels. Difference between overall performance and performance in all other panels was evaluated using a paired Student’s t-test. For characteristic comparison, only variants that affect genes in either better or worse performing panels were considered. Better and worse performance was defined as P value<0.05 and an estimated difference in AUROC above 0.01 and below -0.01, respectively.

## RESULTS

### Evaluation dataset

After collecting all ClinVar variants with a classification of P/LP/B/LB that are annotated in dbNSFP and excluding variants without any assertion criteria provided, a total of 238,977 variants were collected. 112,536 (47%) are classified as B/LB and 126,441 (53%) as P/LP. The majority of the variants (63.1%) were created at or after 2020 and 81,283 (34%) during or after 2023. 46,013 (19.3%), 50,916 (21.3%) and 3,349 (1.4%) of variants were classified as affecting genes with a known AD, AR and XL mode of inheritance, respectively.

### Overall performance

Overall performance was evaluated on several subsets of the data according to the year the variants were created in ClinVar. For each year between 2021 and 2024, all the variants created at or after that year were joined into an evaluation set, each subsequent year is a subset of the previous one. Additionally, for every year, performance was measured for both the entire set of scored variants for each tool (i.e the full set), and a subset of variants that have a score for all of the tools (i.e the complete set) (Table 1). MetaRNN, ClinPred and BayesDel outperformed all other tools across different years and for both the complete and full datasets (Figure 1A) with a AUROC in the complete set of variants created at or after 2023 of 0.984 [0.983,0.985], 0.982 [0.981,0.984] and 0.982 [0.981,0.983], respectively. Setting a threshold corresponding to a precision level of 0.9 (90% probability that a positive variant is pathogenic), all three tools demonstrated high recall values, identifying 83.8%, 81.8% and 79.3% of pathogenic variants, respectively (Table 1). Additional top performing tools include GMVP, REVEL, VARITY, AlphaMissense, VEST4, DEOGEN2, CADD, PrimateAI3d and MetaSVM. Overall performance for these tools remained consistent across years and in both full and complete datasets with minor differences in each tool’s overall performance rank. Reviewing each tool separately, a clear decrease in AUROC was noted for variants created at or after 2020 in four tools, VEST4, ClinPred, MetaRNN and PrimateAI (Table 2). For tools that utilize ClinVar data during training, we tested performance on subsets before and after training date. Out of these tools, only ClinPred and MetaRNN demonstrated a significant decrease in performance for variants created after the training year (−0.189; p=0.004 and -0.158; p=0.008, respectively) (Figure 1B). Overall, for the top performing tools, AUROC was significantly lower for variants created at or after 2020 (−0.011; p<0.0001). We note that despite the decrease in performance, MetaRNN and ClinPred remained among the top performing tools for variants created beyond 2020 and across creation dates. Finally, probability thresholds were calculated for each of the 13 top performing tools using the dataset of variants created at or after 2023, corresponding to the genome-wide calibration. Using a sensitivity cut-off of 0.5% (at least 0.5% of the pathogenic variants are identified using the score threshold), all the tools had scores corresponding to the strong evidence level except one (AlphaMissense), and a single tool (CADD) reached the very strong evidence level (Table 3; Figure 2).

**Figure 1:**
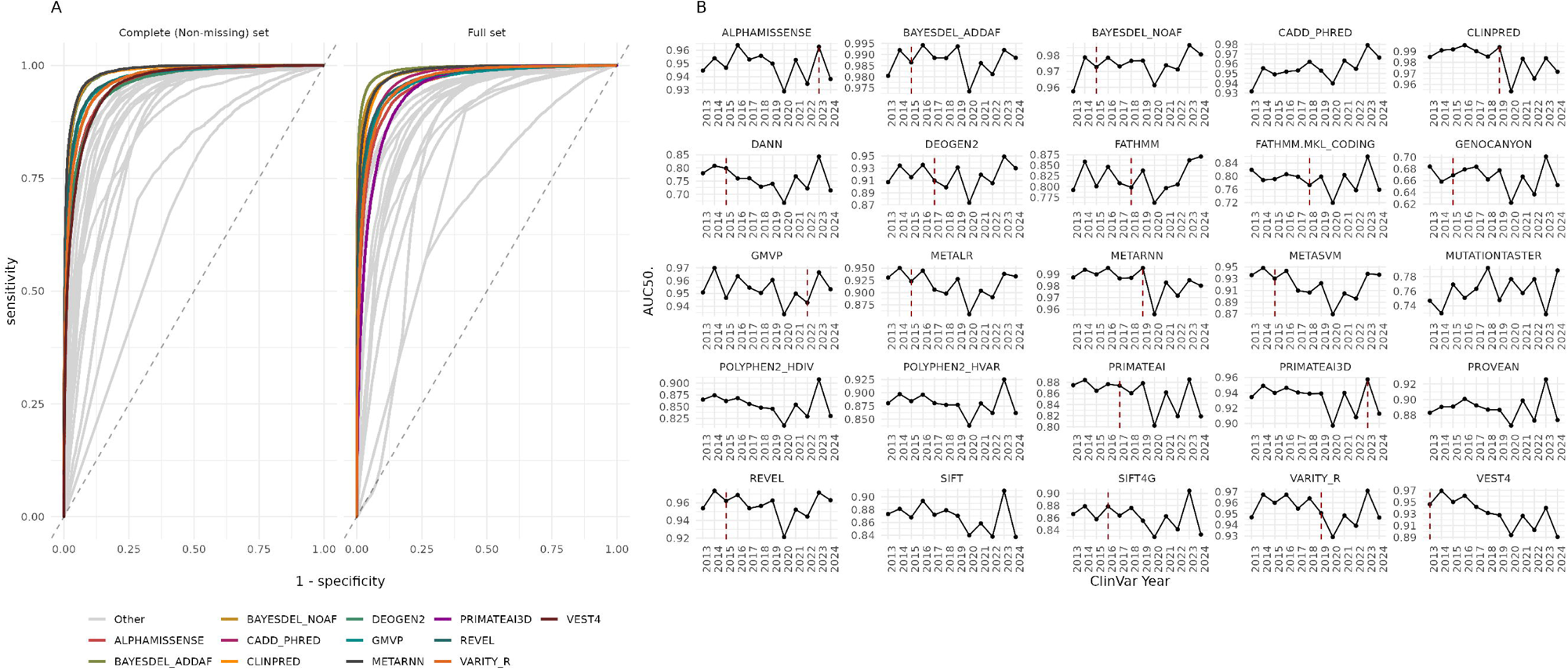
(A) Overall discriminatory performance - ROCs for the different tools on the complete (no missing) and full evaluation sets derived from the list of ClinVar variants added after 01/01/2023. Top performing tools in each subset are colored. (B) Performance by year created - AUROC values calculated for subsets including all the variants created in each year to ClinVar. If a tool was published after 2013, a vertical red line marks the tool’s publication year.

**Figure 2:**
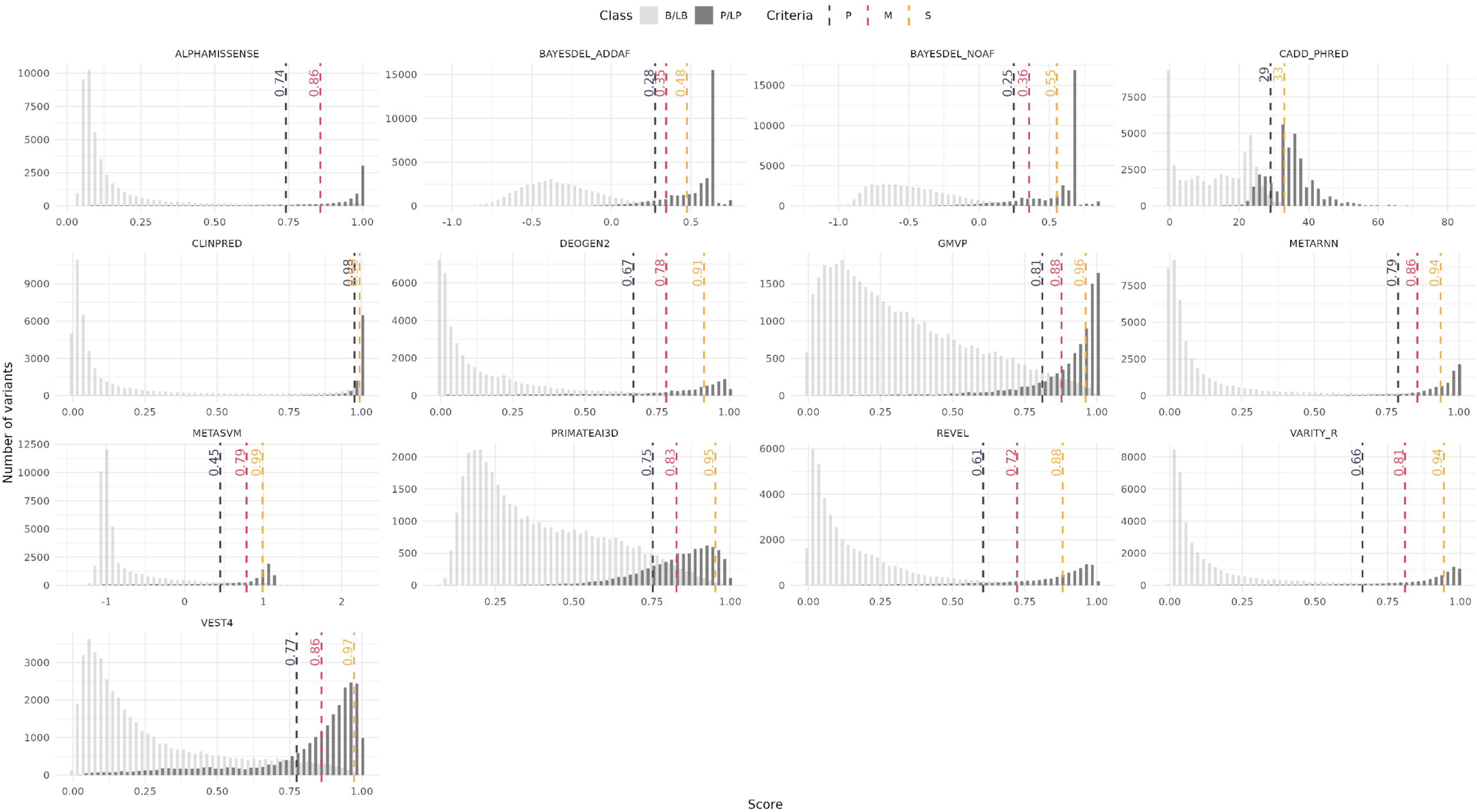
Distribution of scores for P/LP (dark gray) and B/LB (light gray) variants. Recalibrated score thresholds corresponding to various ACMG levels of evidence are added for each score.

### Data missingness

Review of the data revealed varying rates of missing values across different scores in the dbNSFP4.5a database (median 0.371 IQR[0.162,0.404]) (Supplementary Figure 1). PolyPhen2 (0.43), VARITY and PrimateAI3d (0.43) had the highest missingness rate. In contrast, MutationTaster, CADD, DANN, fathmmMKL, Genocanyon, and BayesDel all had a missing rate lower than 0.03. Testing missing rates across the years, we saw a minor, yet significant, increase in missingness throughout the years (0.7% per year; P<0.0001). We note that among the top performing tools, BayesDel and CADD had the lowest missingness rate.

### Conservation

Comparing performance on variants found in regions with various conservation levels (low, intermediate and high) a significant decrease in AUROC was noted as conservation levels increased, with the AUC decreasing by 0.0352 (p=0.0147) and 0.0742 (p<0.0001) in intermediate and high conservation levels, respectively, compared to low levels (Figure 3A). When we reviewed classifications in tools with categorical pathogenicity predictions sensitivity showed a significant increase with higher conservation levels, with an increase of 0.1 (p=0.0001) and 0.199 (p<0.0001) at intermediate and high conservation levels, respectively. The decrease in discriminatory capabilities was mainly due to a major drop in specificity in higher conservation levels with a decrease of 0.191 (p < 0.0001) and 0.43 (p < 0.0001) at intermediate and high conservation levels, respectively, representing a high false positive rate at these conservation levels. Sensitivity and specificity were also high and consistent, indicating robust performance regardless of the conservation context. Reviewing the top performing tools, the effect of conservation level on sensitivity and specificity remained consistent across tools. The overall effect on discriminatory performance (AUROC) was milder compared to all the tools with a decrease of 0.02 (p=0.072) and 0.045 (p<0.0001) in intermediate and high conservation levels, respectively. Comparing the tools on the same set of variants, discriminatory performance was contingent on conservation level, with each level favoring different tools (Figure 3C). Demonstrating these changes, while VEST4 and PrimateAI3d were in the top ten performing tools for variants affecting regions with low conservation, performance of these tools decreased significantly as conservation increased and were no longer present in the list of top performing tools in the high conservation subset. Some tools, such as GMVP and MetaSVM on the other hand increased their overall rank across tools as conservation increased. We then set out to assess the different tools’ performance in varying levels of mismatch constraint. Mismatch constraint refers to the depletion of missense variants in a genomic region compared to what would be expected by chance. Similarly to conservation level, this metric helps identify regions that are intolerant to variation. Overall, a direct association was observed between the mismatch constraint level and discriminatory performance (0.036 increase in AUROC for every 0.1 increase in observed/expected mismatch ratio; p<0.0001) corresponding to better performance in less conserved regions (Figure 3D). Notably, among the top performing tools, VEST4 demonstrated the most significant decrease in performance for the least constrained regions (AUROC=0.891 vs mean AUROC of 0.925), while MetaSVM’s performance decreased in the most constrained regions (AUROC=0.902 vs mean AUROC of 0.934).

**Figure 3:**
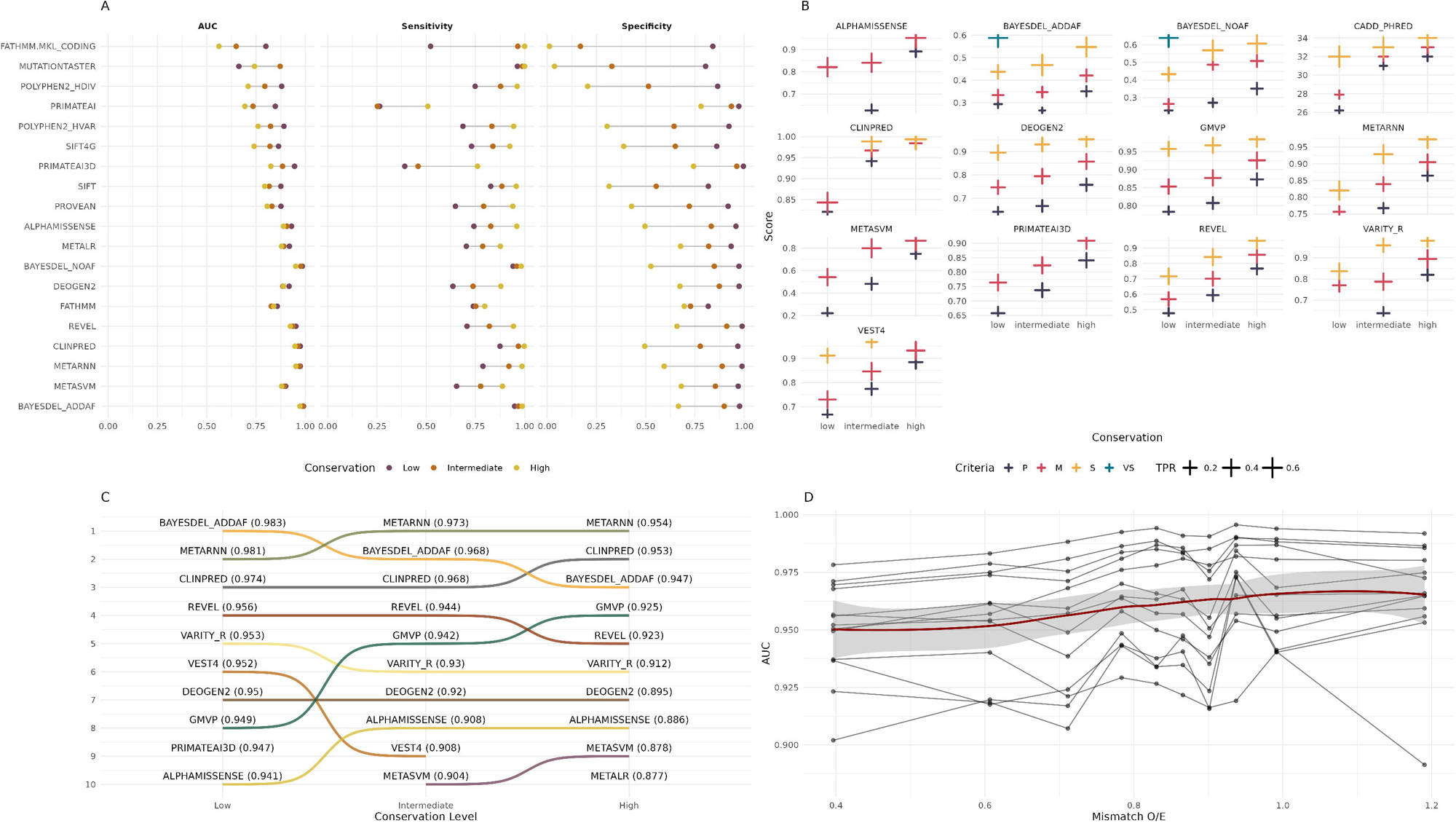
The effect of conservation on performance and calibration. (A) Discriminatory performance metrics (AUC/Sensitivity/Specificity) in each level of conservation. (B) Recalibrated score thresholds corresponding to various ACMG levels of evidence are added for each score in each conservation level. (C) Bump plot demonstrating the change in rank by conservation level for the top performing tools ranked by their AUC levels. (D) AUC for each tool in various levels of mismatch O/E rates, demonstrating a decrease in performance as conservation increases and mismatch O/E decreases.

### Allele Frequency

Variant allele frequency has been utilized during the development of various VEP, either as a feature^18,24,25^, or in order to define which variants may be categorized as benign when generating the training set^22,27^. These allele frequencies may be significantly different among diverse populations. Therefore, we set out to evaluate how allele frequency affects VEP performance. Comparing performance between each allele frequency category and the AF=0 category (variant not found in gnomAD), we see a significant decrease in performance. This observation was seen both when considering all the VEP tools (P<0.01) and the top performing ones (P<0.05). Decrease in performance was most significant in variants with AF>10^−4^ (estimated AUC decrease = -0.032; P<0.0001). When evaluating how AF affects each tool’s performance, five tools demonstrated a significant decline in AUROC as AF increases (CADD_PHRED, BAYESDEL_NOAF, VARITY_R, BAYESDEL_ADDAF and VEST4) and a single tool (CLINPRED) demonstrated a significant increase in AUROC as AF increased (Figure 4A). Recalibrating scores for each AF category, we note that while the version of BayesDel which does not incorporate AF as a feature (BAYESDEL_NOAF) did not demonstrate difference in the scores required to reach the “Strong” level of evidence, the version in which AF is an added feature (BAYESDEL_AF), demonstrated a decrease in score thresholds as AF increased, corresponding to lower overall scores given to more common variants (Figure 4B; Table 4).

**Figure 4:**
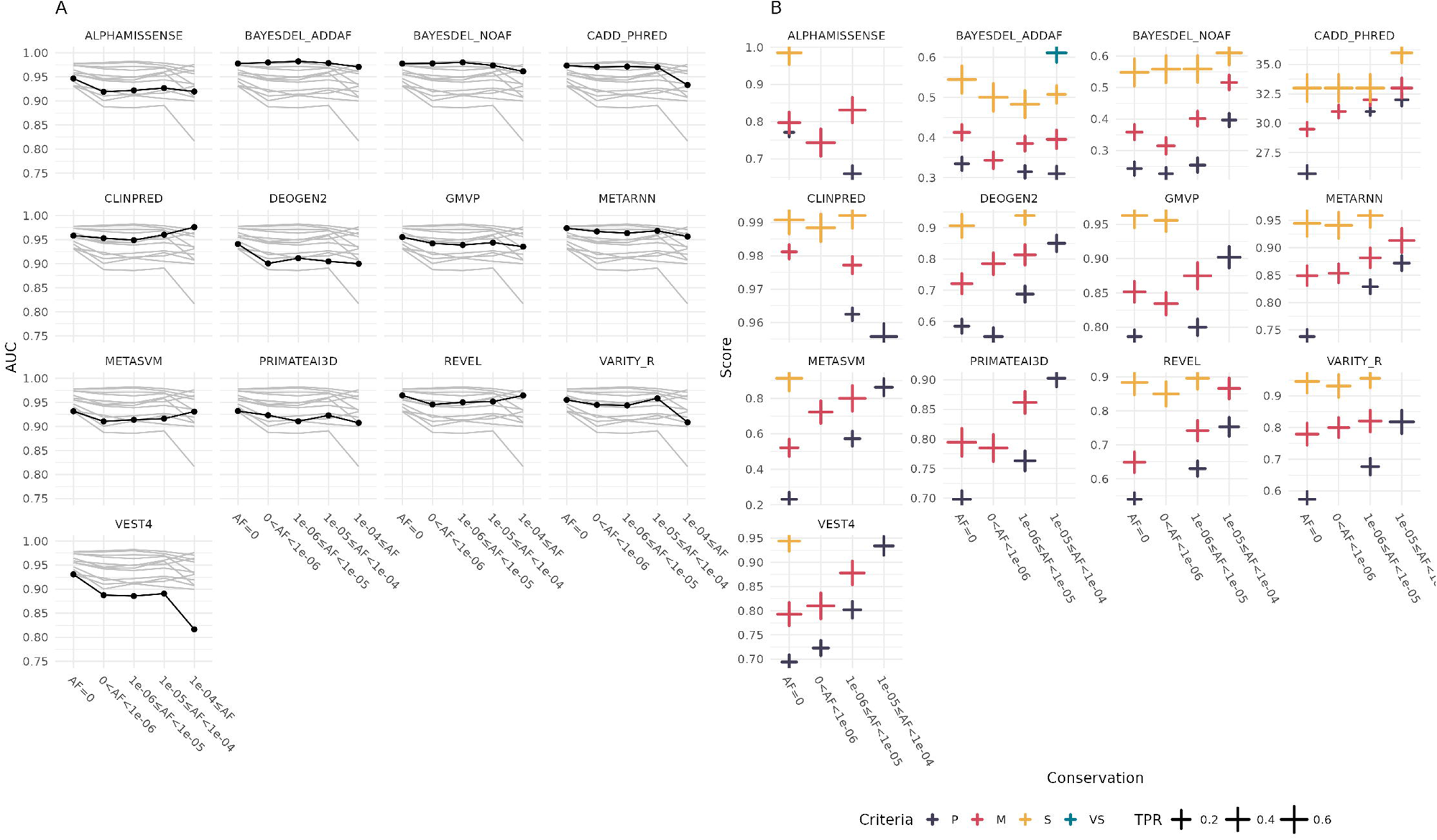
Performance stratified by allele frequency (AF). (A) AUC for each tool in increasing levels of AF. (B) Recalibrated score thresholds corresponding to various ACMG levels of evidence are added for each score in each AF rate.

### Mode of inheritance

In order to evaluate how mode of inheritance (MOI) affects prediction tool performance, variants were categorized according to whether they affect AD, AR and XL genes. To complement this evaluation, pLI (probability of loss-of-function intolerance) and pRec scores were also collected for each gene, and each gene was marked as pLI high (pLI>0.9) and pRec high (pRec>0.9). pLI is a metric used to identify genes that are intolerant to loss-of-function (LOF) variation^3^. pRec is defined as the probability of being intolerant of homozygous, but not heterozygous LoF variants. Overall, most tools (19/25) performed better on AR variants compared to AD (Figure 5A) (0.025 difference in AUROC; p=0.032). The majority of the tools also performed better on pRec variants but the difference in performance compared to pLI high was not significant (p=0.086).

**Figure 5:**
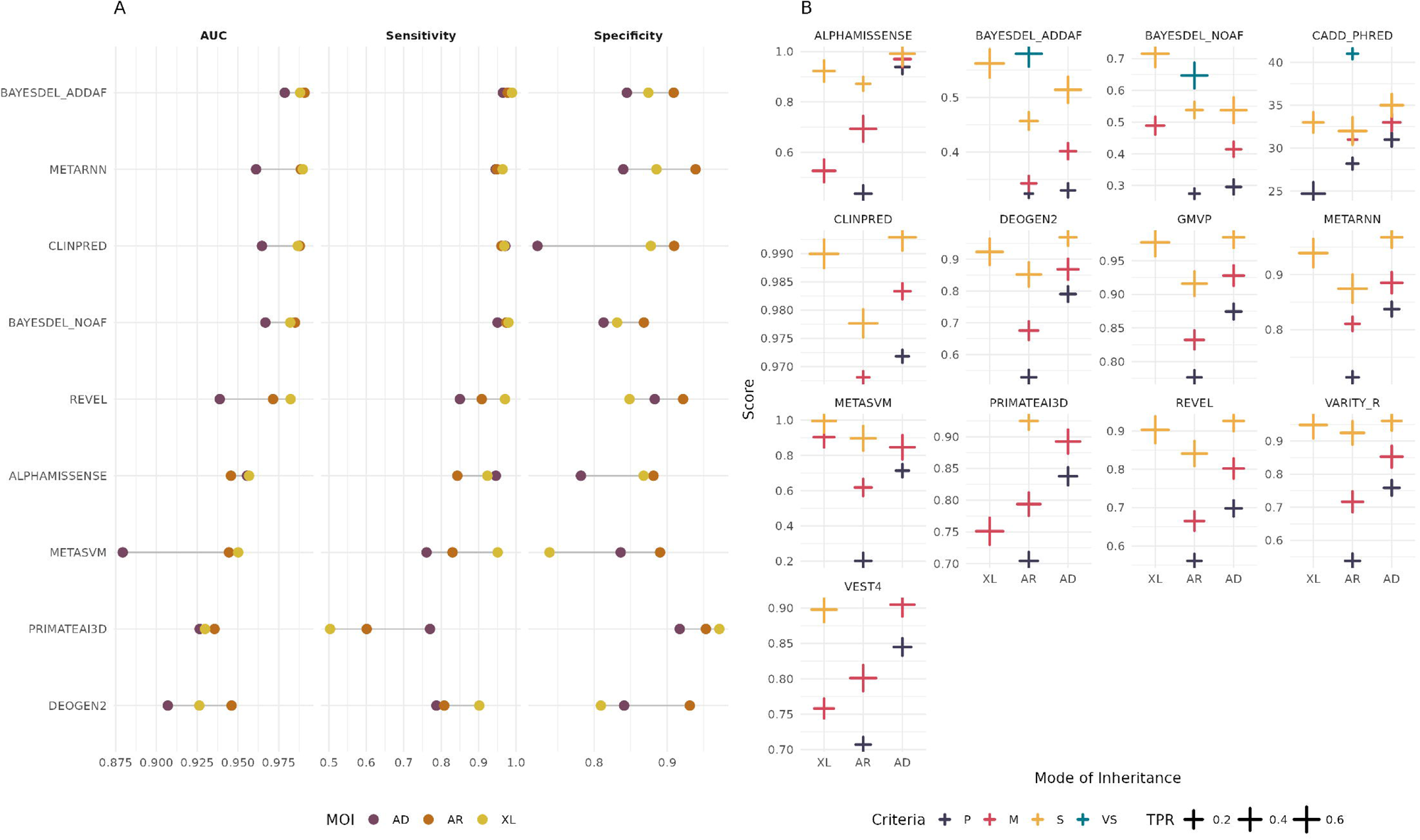
Performance stratified by mode of inheritance (MOI), into autosomal dominant (AD), autosomal recessive (AR) and X-linked (XL). (A) Discriminatory performance metrics (AUC/Sensitivity/Specificity) in each MOI. (B) Recalibrated score thresholds corresponding to various ACMG levels of evidence are added for each score in each MOI.

Out of the top performing tools, AlphaMissense and VEST4 were the only tools performing better on AD variants. Recalibration of scores corresponding to the different levels of evidence, demonstrated that for the majority of the tools, criteria thresholds may be lowered when classifying variants affecting genes associated with an AR MOI compared to AD. For example, reviewing re-calibrated thresholds for REVEL, while in AD variants, we observe similar thresholds compared to the ones suggested by the SVI (0.698, 0.802 and 0.926 for supporting, moderate and strong evidence levels, respectively), in AR genes the thresholds were significantly lower (0.561, 0.665, 0.841) (Table 5; FIgure 5B).

### Disease category

Targeted sequencing of gene panels utilizes pre-generated gene panels, composed of genes with putative association with specific phenotypes in order to reduce costs and enhance downstream analysis. Given a set of established, expert-curated gene panels, we reviewed whether VEP performance is significantly altered in certain panels. First, overall performance for the top performing tools was assessed for each gene panel. Out of 62 panels, performance was significantly better in 11 (17.7%) panels and significantly worse in 9 (14.5%) (Figure 6). Specifically, 55% of the panels with worse performance were associated with hematologic or solid malignancy. Association with inflammatory disorders (3/11), connective tissue disorders (2/11) and neurologic disorders (3/11), was observed among panels with improved performance. Comparing variant characteristics in panels associated with better or worse performance, we saw a significantly higher rate of XL variants in panels with better performance (3.5% vs 0.1%; P<0.0001). We also noted that the rate of pathogenic variants in regions with low conservation was significantly higher in panels with worse performance compared to those with better performance (45.7% vs 28.4%; P<0.0001).

**Figure 6:**
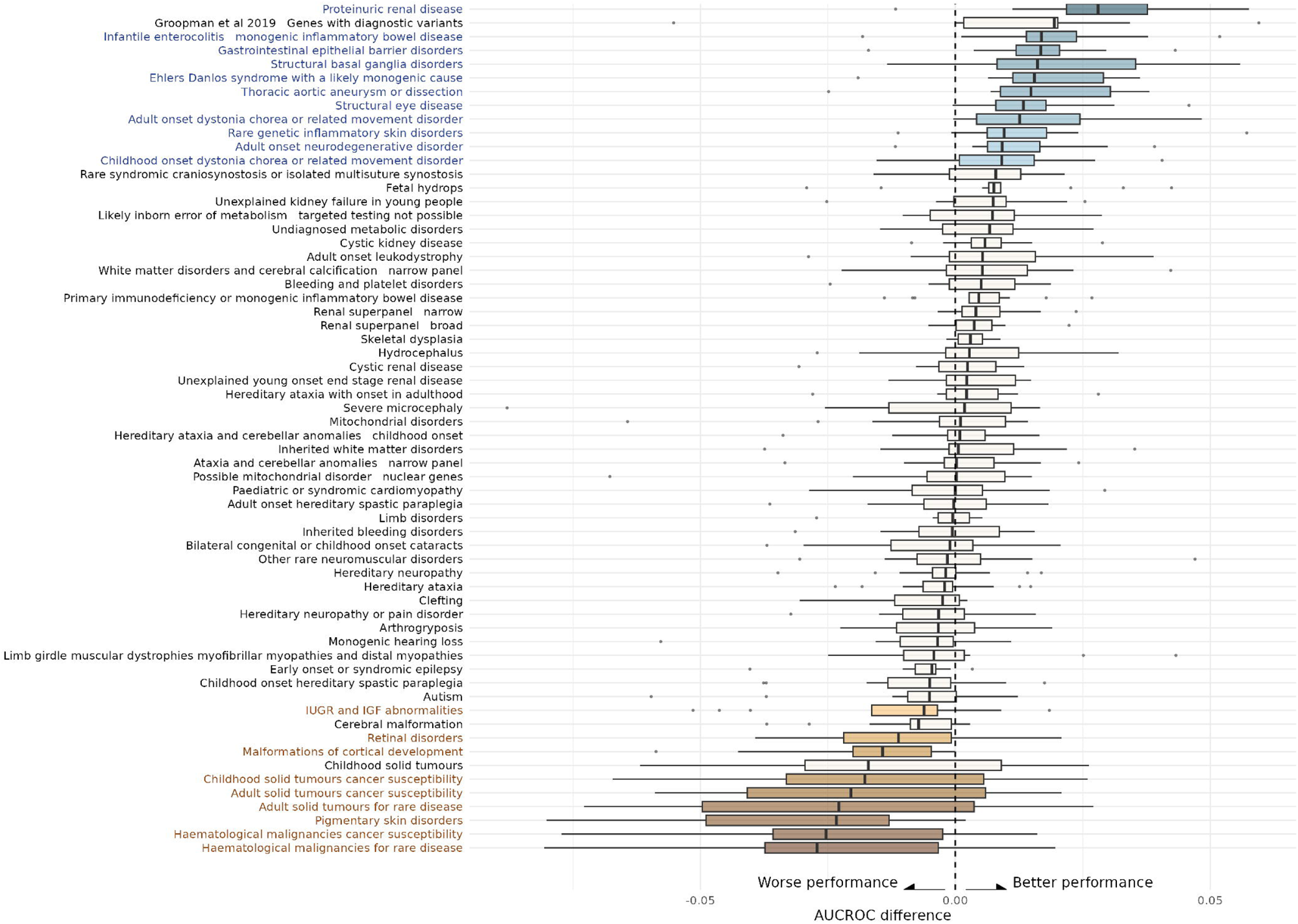
Performance differences across disease categories. For each tool, AUROC was calculated for each disease group. AUROC difference was calculated between each disease category and all other disease categories. The plot demonstrates the distribution of these differences across disease categories. Disease categories with a significant improvement or reduction in AUROC were marked in blue and brown, respectively.

## DISCUSSION

In this study, we test the performance of various VEP tools under different conditions and demonstrate significant variability in discriminatory performance. We also calculate, for each tool, under each condition, score thresholds corresponding to the various levels of evidence for variant classification purposes. We show that even in conditions where the discriminatory performance remains high, score distributions may shift and thresholds must therefore be recalibrated in order to correspond to the correct level of evidence.

### Overall performance, data missingness and variant creation dates

Initially, we review missing values for each tool, demonstrating a large inter-tool discrepancy in score availability in the dbNSFP database. Since this database is commonly used for variant annotations and classification, it is important to select tools with a minimal rate of missing values in order to ensure consistency and refraining from switching between tools during variant classification. We then review how different tools perform on subsets stratified by the variant creation date. We identify a mild yet significant decrease in performance for variants that were created after 01/2020 most probably due to the higher rate of variants used during training of many of the tools in earlier years. We therefore selected a subset of the data corresponding to newly created variants, not present in the database before 2023, for the downstream analysis. Using a subset of this dataset, which includes only variants with scores for every tool, we identify the tools with the best performance genome-wide.

### Performance under different scenarios

In the field of machine learning, the term ‘fairness’ refers to the ability of prediction models to provide consistent and unbiased results across different subsets of data. In variant pathogenicity prediction, fairness entails the model’s consistent performance across various variant subsets, stratified by characteristics such as conservation, MOI allele frequency and disease association. We evaluated how conservation affects performance, with the majority of VEP tools demonstrating a decrease in performance in highly conserved regions compared to lower conservation levels. The decrease in performance was mainly due to a decrease in specificity in highly conserved regions. In these regions, benign variants may still receive high pathogenicity scores simply because they occur in conserved areas therefore leading to a higher rate of false positives. We also evaluate how MOI affects performance, and similarly, we show that performance is significantly better for variants affecting AR and XL genes compared to AD genes. Gerasimavicius et al. analyzed how putative molecular mechanisms affect the performance of various VEP. In their study they show that nearly all variant effect predictors perform worse at identifying non-loss-of-function mutations compared to loss-of-function mutations and on AD variants compared to AR. AR disorders often result from loss-of-function variants that completely disrupt gene function. Such variants tend to have more pronounced molecular effects and are therefore more easily identified by VEPs. Interestingly, among the top performing tools, only AlphaMissense demonstrated better discriminatory performance in AD variants compared to AR. We attribute this difference to AlphaMissense’s training method, in which a benign state was defined using an AF threshold. As AR variants tend to have higher AF, utilizing AF for this purpose may incorporate bias into the training set, marking P/LP AR variants as benign. When comparing performance across disease categories, we noted a significant decrease in performance in genes associated with solid and hematologic malignancies. We attribute this difference to an increase in pathogenic variants in regions of low conservation, possibly because mutations there are more tolerable to cells and can contribute to cancer development without causing cell death. As our work demonstrates, VEP tools assign lower scores for variants found in low conservation regions and therefore are more likely to miss pathogenic variants in these regions. Another reason for a decrease in performance in malignancy associated genes is the fact that most hereditary cancer syndromes are transmitted by AD mechanisms^34^, in which our work demonstrates, most tools perform worse.

### Score threshold re-calibration

In 2022, ClinGen SVI issued a revised guideline pertaining to the use of the PP3/BP4 criteria^4^. In this work, VEP performance was recalibrated using a genome-wide data set and thresholds corresponding to different levels of evidence were determined. Our study joins previous descriptions of variability in VEP performance under different molecular and clinical scenarios^7^. It is therefore unlikely that the thresholds determined by the SVI are generalizable to every possible variant. In our study, we recalibrated these thresholds, for the top performing tools under different scenarios. Recalibration according to conservation levels suggested that score thresholds should be lowered when classifying variants in regions with low conservation (Figure 3B; Table 6). Since conservation level is incorporated in most VEP tools, variants in highly conserved regions often receive higher pathogenicity scores due to the assumption that any change in a conserved region is more likely to be deleterious. Therefore, in these regions, higher thresholds are needed to distinguish truly pathogenic variants from benign ones that also receive high scores (Supplementary figure 1).

Recalibration according to MOI suggested a decrease in score thresholds when classifying AR variants compared to AD variants, and in some tools, improved evidence stratification reaching the strongest level of evidence. A possible reason for lower scores corresponding to P/LP AR variants could be the underlying characteristics of such variants, including lower conservation and higher allele frequencies, which drive prediction scores lower, compared to AD variants. Regardless of the reason, our analysis demonstrates that while overall discriminatory performance remains high in the variant subsets tests, the distribution of scores differs significantly and therefore require modifications in score thresholds in order to more accuratel correspond to the various levels of evidence.

### Limitations

Our analysis is derived from the ClinVar database. In order to reduce the likelihood of testing a tool’s performance on variants already used during its training, we selected only variants created after 01/2023. It is possible that some of the VEP tools tested in this study have been used to classify variants in our evaluation test, which might introduce bias into overall performance assessments. This bias may affect overall performance assessment, but is not expected to alter the study’s findings regarding altered performance under different scenarios.

In our study, we utilize a non-parametric density function to estimate likelihoods. A crucial aspect of this method is selecting an appropriate bandwidth for smoothing the data. The bandwidth determines how much the data is smoothed: a smaller bandwidth captures more details but may introduce noise, while a larger bandwidth smooths out noise but may overlook important features. We selected Silverman’s ‘rule of thumb’ as our bandwidth smoothing method^35^, and found that it offered an acceptable trade-off between accurately representing the true distribution of scores and minimizing the noise introduced by over-granularity. This allowed us to generate reliable likelihood estimates that are both accurate and robust against random fluctuations in the data.

## Conclusion

By recalibrating thresholds based on different molecular and clincial scenarios, the tools can more accurately stratify variants according to their true pathogenic potential. This refinement enhances the confidence in variant classification, allowing some variants to reach the strongest level of evidence for pathogenicity as per the ClinGen SVI framework. This study emphasizes the importance of analyzing VEP tools in different clinically relevant contexts in order to more accurately apply them in clinical settings.

## Supporting information

Manuscript tables

## ABBREVIATIONS

AD: Autosomal dominant
AR: Autosomal recessive
AUROC: Area under the ROC curve
B/LB: benign / likely benign
LOF: Loss of function
MOI: Mode of inheritance
P/LP: pathogenic / likely pathogenic
pLI: probability of loss-of-function intolerance
ROC: Receiver operating characteristic
SVI: ClinGen Sequence Variant Interpretation working group

---

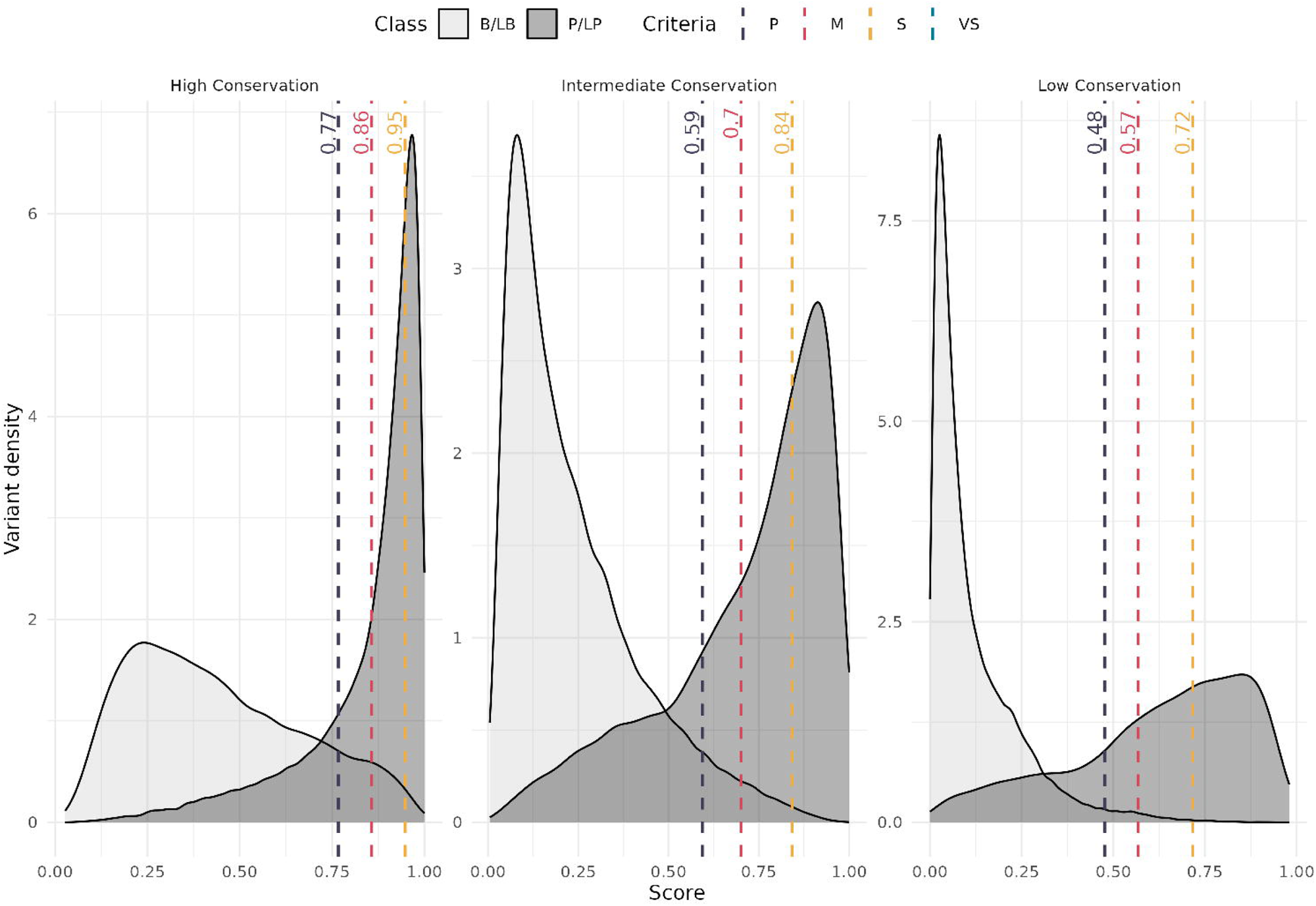

